# Aceclofenac Fast Dispersible Tablet Formulations: Effect of different concentration levels of Avicel PH102 on the compactional, mechanical and drug release characteristics

**DOI:** 10.1101/773671

**Authors:** Riffat Yasmin, Muhammad Harris Shoaib, Farrukh Rafiq Ahmed, Faaiza Qazi, Huma Ali, Farya Zafar

## Abstract

Objective of this study was based on the formulation development of fast dispersible Aceclofenac tablets (100mg) and to evaluate the influence of pharmaceutical mixtures of directly compressible Avicel PH102 with Mannitol and Acdisol on the compressional, mechanical characteristics and drug release properties. Fifteen different aceclofenac formulations were developed by central composite rotatable design (CCRD). Among them best possible formulations (FA–FH) were selected on the basis of micromeritic properties, appropriate tablet weight and disintegration time for further study. Tablets were compressed by direct compression method using hand held hydraulic press with a compressional force ranging from 8 to 80 MN/m^2^ (MPa). Pre and post compression studies were performed and the compressed formulations (FA-FH) were assessed for different quality tests. The Heckel and Kawakita equations were applied for determination of compressional behavior of formulations. The quality attributes suggested that formulation (FB) containing avicel PH 102 (20%), mannitol (25%) and ac-di-sol (3%) as best optimized formulation showing better mechanical strength i.e. hardness 37.75 ± 0.14N, tensile strength 5.67MN/m^2^ and friability 0.34%. Furthermore, compressional analysis of FB showed lowest P_Y_ value 59.52 MN/m^2^ and P_k_ value 1.040 MN/m indicating plasticity of the material. Formulation FB disintegrated rapidly within 21 seconds and released 99.92 % drug after 45min in phosphate buffer pH 6.8. Results of drug release kinetics showed that formulations FA-FH followed Weibull and First-order models in three different dissolution media. Avicel based formulation mixture exhibit excellent compactional strength with rapid disintegration and drug release.

## INTRODUCTION

Advancement in tablet manufacturing technology has offered viable dosage alternatives for those patients who are facing problem related to compliance with conventional dosage forms. One such alternative dosage form is the fast dispersible tablets Fukami et al. (*1*). These tablets are of two types: first type is taken in mouth without water to disintegrate rapidly or disperse readily and the second type of tablets form dispersion or solution in water to be taken by patients Schiermeier and Schmidt (*2*, Martin et al. (*3*). These tablets are usually developed by direct compression method. Aceclofenac is a Cycloxygenase inhibitor having analgesic and anti-inflammatory activity. Due to its short half-life (4hr) and twice daily dose, it is considered as suitable candidate for fast dispersible tablets Setty et al. (*4*, Parfitt (*5*).

Avicel^®^, the first commercialized brand of microcrystalline cellulose (MCC), is introduced by FMC Corporation as a direct compression tableting ingredient. MCC is a partially depolymerized cellulose that is obtained as a pulp by mineral acid treatment of alpha cellulose type lb of fibrous plant material. Cellulose is the most abundant natural polymer having linear chains of b-1, 4-D anhydroglucopyranosyl units. Pharmaceutical MCC is most commonly obtained from wood where cellulose chains are packed in layers held together by strong hydrogen bonds and lignin (cross-linking polymer). The primary particles of all MCC types (101, 102 and 200) are about 50µm but difference in the larger aggregated particle numbers. Type 102 have a median particle size of about 100µm indicating adequate flow properties for successful tableting. MCC deforms plastically and maximizes the interparticle bonding area during compression. It forms strong and cohesive compacts even under low compression pressure due to formation of numerous hydrogen bonds. Tableting is further enhanced by mechanical interlocking of elongated and irregularly shaped particles. (*6*).

During tablet manufacturing, compression of powder shows reduction in volume of powder bed by the application of compressional pressure in a confined space. As a result, there is formation of strong inter-particle bonds which produce a compact mass and built inherent strength in the compact to increase the overall mechanical strength of the dosage unit. This analysis is significant in understanding the behavior of poorly compactable powders during the manufacturing of the tablets Nicklasson (*7*).

### Heckel analysis

It is one of the most widely used method to relate the reduction of powder bed volume upon application of compressional force. It is based on an assumption that “compression of powder follows first-order kinetics in which pores are reactant and powder densification is the product”. It is demonstrated as follows:

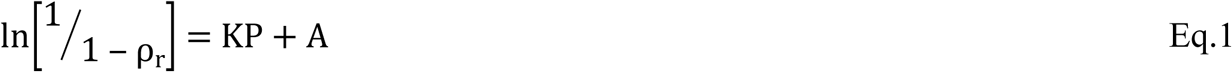

Graph between densification i.e. 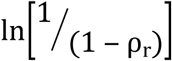 and applied pressure ‘P’ provides the value of slope ‘K’ and intercept ‘A’ Eckert et al. (*8*). Powder densification takes place in three phases i.e. D_o_, D_A,_ and D_B_. D_o_ (relative density) shows the powder densification at die filling stage. Since it measures the packing characteristics of powder, thus high value of D_o_ is an indication of highly dense packing of powder blend Mahmoodi et al. (*9*, Denny (*10*). The D_B_ value indicates densification of powder when it is compressed and particles show movement and re-arrangement. Magnitude of rearrangement based on the theoretical point of densification at which deformation of particles started Adetunji et al. (*11*).

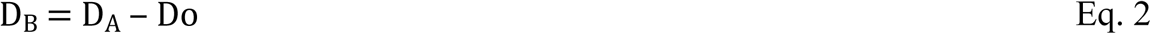

The D_A_ value is the final compact densification (D_A_= ρ_r_) and it is calculated from intercept ‘A’ of the Heckel plot:

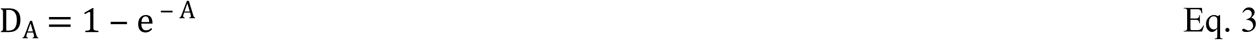

### Kawakita analysis

Kawakita equation describes the compressional behavior of powder, either by tapping or from continuous compression experiment Kawakita and Lüdde (*12*). It explains the relationship between degree of volume reduction ‘C’ upon the application of compressional pressure ‘P’. The linear expression of Kawakita equation is given below:

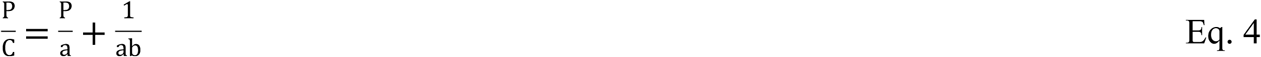

Or

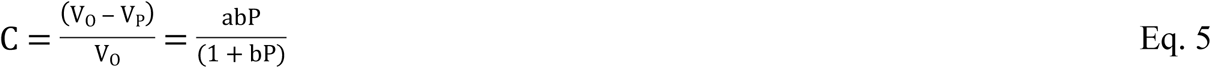

Where, V_O_ is the initial bulk volume of the powder, V_P_ is the volume of the powder after compression, ‘a’ is the total volume reduction for the powder bed or minimum porosity before compression and ‘b’ is the powder’s plasticity Adetunji and colleagues (*11*).

The aim of the present study was to prepare fast dispersible Aceclofenac tablets and to evaluate the compressional behavior of the newly developed tablets under recommended condition of tablet manufacturing using Heckel and Kawakita analysis. Formulations were designed by Central Composite Design (CCD) using varying concentration of avicel PH102, mannitol and ac-di-sol. All the formulations were developed by direct compression method.

## MATERIALS

Aceclofenac was gifted by Sami Pharmaceutical (Pvt.) Limited. Avicel PH-102, mannitol, ac-di-sol, aspartame, talc and vanilla flavor were purchased from FMC Corporation, USA.

## METHODS

### Experimental Design

Twenty fast dispersible Aceclofenac tablet formulations (100mg) were designed with the help of Central Composite Design using Design Expert®10.0 software (Stat –Ease, Inc, Minneapolis, MN 55413, USA). The selected independent variables were avicel PH102 (20-35%), mannitol (10– 25%) and ac-di-sol (1–3%). Excipients such as aspartame, vanilla flavor and talc were used at a fixed concentration i.e. 2%, 1%, and 2% respectively, composition is given in Table 1. Disintegration time, percentage friability and hardness of formulations were selected as the dependent variables. Response surface methodology (RSM) was used to explore the interaction of avicel PH102, mannitol and ac-di-sol to establish the appropriate amount of excipient for optimized fast dispersible formulation. On the basis of fit summary, ANOVA and multiple correlation coefficient appropriate model was selected.

**Table 1:**
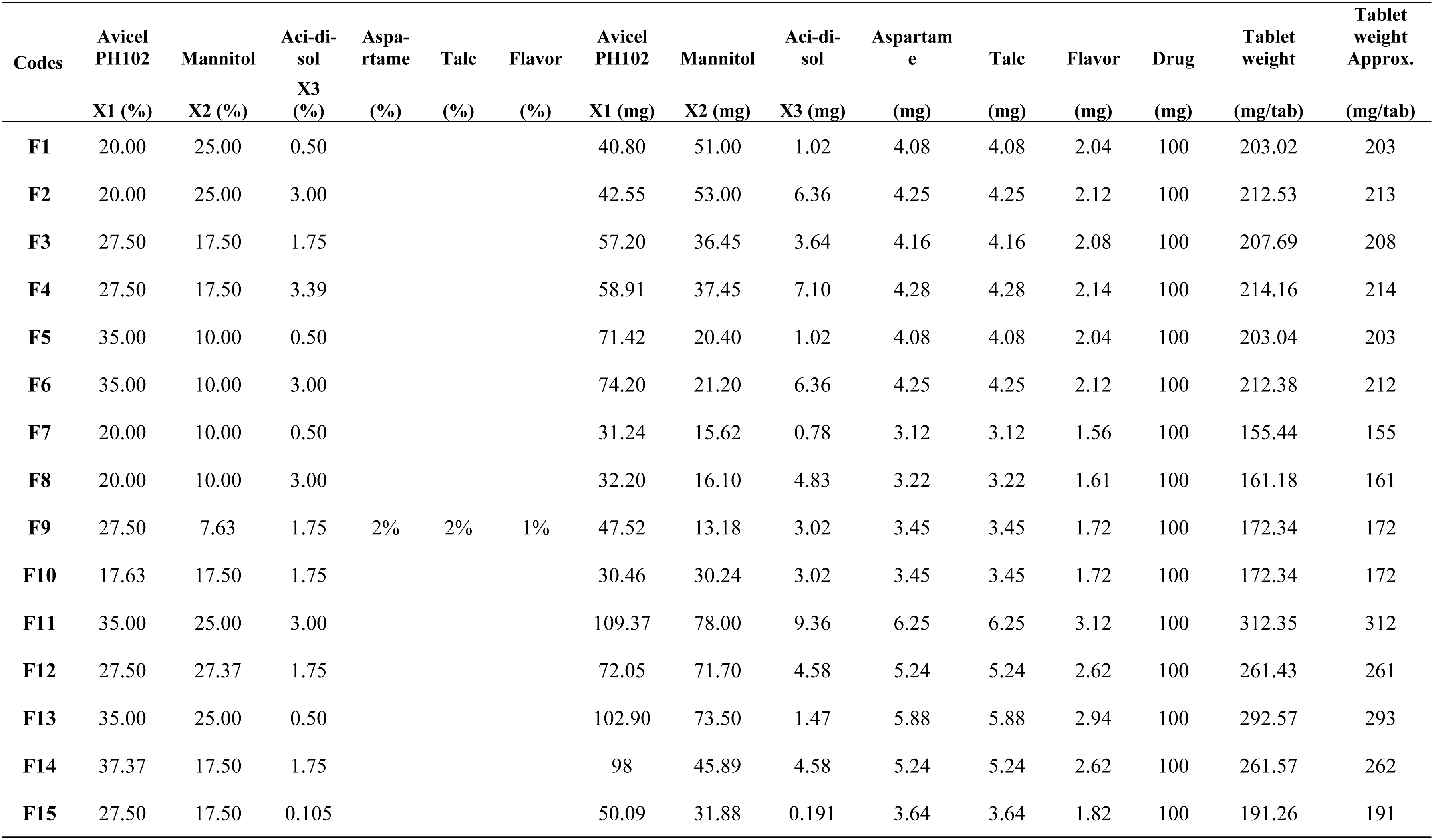
Composition of fast dispersible aceclofenac formulations using central composite design.

| Codes | Avicel<br>PH102 | Mannitol | Aci-di-sol<br>X3<br>(%) | Aspa-<br>rtame<br>(%) | Talc<br>(%) | Flavor<br>(%) | Avicel<br>PH102<br>X1 (mg) | Mannitol<br>X2 (mg) | Aci-di-sol<br>X3 (mg) | Aspartam<br>e<br>(mg) | Talc<br>(mg) | Flavor<br>(mg) | Drug<br>(mg) | Tablet<br>weight<br>(mg/tab) | Tablet<br>weight<br>Approx.<br>(mg/tab) |
| --- | --- | --- | --- | --- | --- | --- | --- | --- | --- | --- | --- | --- | --- | --- | --- |
| **F1** | 20.00 | 25.00 | 0.50 |  |  |  | 40.80 | 51.00 | 1.02 | 4.08 | 4.08 | 2.04 | 100 | 203.02 | 203 |
| **F2** | 20.00 | 25.00 | 3.00 |  |  |  | 42.55 | 53.00 | 6.36 | 4.25 | 4.25 | 2.12 | 100 | 212.53 | 213 |
| **F3** | 27.50 | 17.50 | 1.75 |  |  |  | 57.20 | 36.45 | 3.64 | 4.16 | 4.16 | 2.08 | 100 | 207.69 | 208 |
| **F4** | 27.50 | 17.50 | 3.39 |  |  |  | 58.91 | 37.45 | 7.10 | 4.28 | 4.28 | 2.14 | 100 | 214.16 | 214 |
| **F5** | 35.00 | 10.00 | 0.50 |  |  |  | 71.42 | 20.40 | 1.02 | 4.08 | 4.08 | 2.04 | 100 | 203.04 | 203 |
| **F6** | 35.00 | 10.00 | 3.00 |  |  |  | 74.20 | 21.20 | 6.36 | 4.25 | 4.25 | 2.12 | 100 | 212.38 | 212 |
| **F7** | 20.00 | 10.00 | 0.50 |  |  |  | 31.24 | 15.62 | 0.78 | 3.12 | 3.12 | 1.56 | 100 | 155.44 | 155 |
| **F8** | 20.00 | 10.00 | 3.00 |  |  |  | 32.20 | 16.10 | 4.83 | 3.22 | 3.22 | 1.61 | 100 | 161.18 | 161 |
| **F9** | 27.50 | 7.63 | 1.75 | 2% | 2% | 1% | 47.52 | 13.18 | 3.02 | 3.45 | 3.45 | 1.72 | 100 | 172.34 | 172 |
| **F10** | 17.63 | 17.50 | 1.75 |  |  |  | 30.46 | 30.24 | 3.02 | 3.45 | 3.45 | 1.72 | 100 | 172.34 | 172 |
| **F11** | 35.00 | 25.00 | 3.00 |  |  |  | 109.37 | 78.00 | 9.36 | 6.25 | 6.25 | 3.12 | 100 | 312.35 | 312 |
| **F12** | 27.50 | 27.37 | 1.75 |  |  |  | 72.05 | 71.70 | 4.58 | 5.24 | 5.24 | 2.62 | 100 | 261.43 | 261 |
| **F13** | 35.00 | 25.00 | 0.50 |  |  |  | 102.90 | 73.50 | 1.47 | 5.88 | 5.88 | 2.94 | 100 | 292.57 | 293 |
| **F14** | 37.37 | 17.50 | 1.75 |  |  |  | 98 | 45.89 | 4.58 | 5.24 | 5.24 | 2.62 | 100 | 261.57 | 262 |
| **F15** | 27.50 | 17.50 | 0.105 |  |  |  | 50.09 | 31.88 | 0.191 | 3.64 | 3.64 | 1.82 | 100 | 191.26 | 191 |

#### Pre-Compression Studies

##### Assessment of powder densities and flow Characteristics

Bulk and tapped densities of all formulation blends were determined by pouring powder mixture in a 100 ml measuring cylinder and their initial and tapped volumes were recorded. Powder blends were assessed by Hausner’s Ratio (HR), compressibility index (C.I) and angle of repose (Θ) using following equations Lin and Cham (*13*):

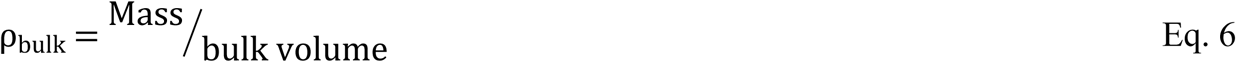

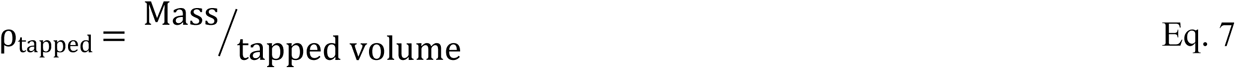

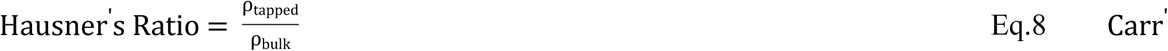

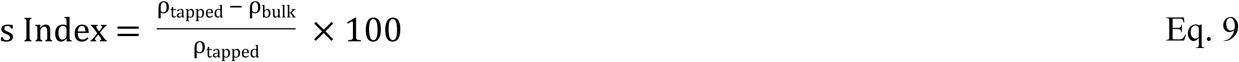

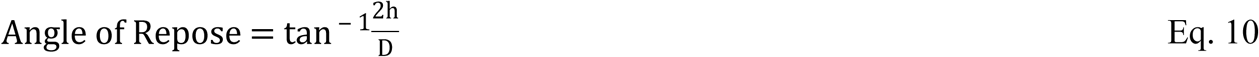

Where ρ_bulk_ and ρ_tapped_are the bulk and tapped densities of powder blends respectively. In the above equation of angle of repose, ‘h’ is the height of powder heap and ‘D’ is the diameter of the heap formed. True density (ρ_t_) of all powder blends were determined by *liquid displacement method* using xylene as displacement liquid with the help of pycnometer. The difference between empty pycnometer (W) and xylene filled pycnometer (W_1_) was calculated as the weight of xylene (W_2_). Approximately 2 g sample of each formulation was weighed (W_3_) and transferred to the pycnometer along with xylene and weighed again (W_4_). True density was calculated using following equation Odeku et al. (*14*):

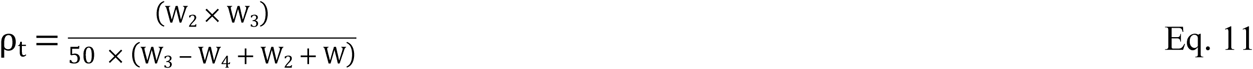

#### Compression of powder blends

All the ingredients were weighed and passed through 20 mesh sieve separately and mixed for six minutes (optimized mixing time) by tumbling in a polybag. After mixing all the ingredients, talc was added and mixed for further five minutes. Blends were compressed by manual filling of the die cavity. Tablets were compressed with weight ranged from 0.155 – 0.212 gm by applying varied compressional pressure 7.72, 23.16, 30.88, 38.0, 46.32, 54.04, 61.76, 69.48 and 77.2 MN/m^2^) using hand held hydraulic tableting machine (locally manufactured) fitted with pressure gauge. Tablets were stored in a desiccator for 24 h over silica gel for elastic recovery and hardening.

#### Evaluation of physicochemical parameters of Aceclofenac formulations

Compressed tablets were weighed and their thickness and diameter were accurately measured with digital vernier caliper (Digital Caliper: Seiko brand). Tablets hardness was determined by using hardness tester (OSK Fujiwara, Ogawa Seiki Co. Ltd., Tokyo, Japan). Tensile strength of round flat faced tablets was calculated using given equation Fell and Newton (*15*):

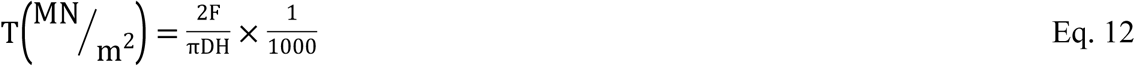

Where, F (N) is the crushing load applied on tablets, ‘H’ is the thickness (cm) and ‘D’ is the diameter (cm) of tablets. Percentage friability was determined using Roche friabilator (H. Jurgens Gmbh H and Co-Bremen, D2800, Germany) with the help of following formula:

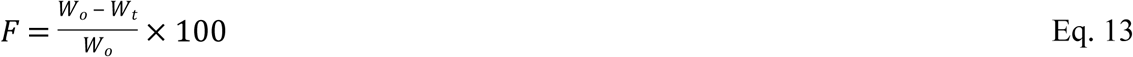

Where W_o_ is the initial weight and W_t_ is the final weight of tablets after 100 rotations. For disintegration test of fast dispersible tablets, 900 mL distilled water was maintained at 37°C. Tablets were disintegrated in seconds Eur.Pharm (*16*). The assay assessment was carried out using UV-Spectrophotometer (UV-1800 Shimadzu Corporation Kyoto, Japan) at 274 nm absorbance Bhardwaj et al. (*17*, Sharma et al. (*18*). Similarly single point dissolution test was performed using phosphate buffer pH 6.8. Samples were collected after 45 minutes and analyzed at 274 nm spectrophotometrically for estimating the percentage drug release Bhardwaj and colleagues (*17*, Sharma and colleagues (*18*).

#### Evaluation of Compressional Behavior

##### Method of applying Heckel equation

Heckel equation is used to relate the powder bed’s relative density (ρ_r_) to the applied pressure during compression which is the ratio of apparent density (ρ_A_) of the tablet and the true density (ρ_T_) of powder blend:

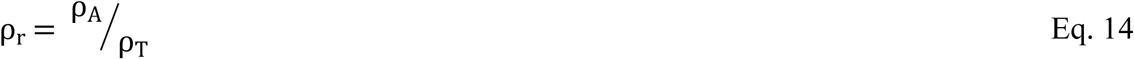

The apparent density ρ_A_ of tablet was calculated using the weight (g), radius (cm) and thickness (cm) of the tablet, whereas true density of the powder blend was determined by liquid displacement method (Eq. 15).

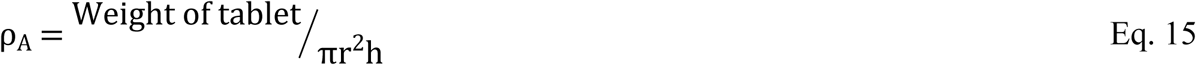

Using the Heckel equation, plot between 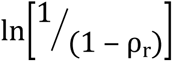 and applied pressure [P] provides the value of K and A. The value of K (slope of the plot) determines the mean yield pressure (1/K= P_Y_) Eckert and colleagues (*8*). This mean yield pressure P_Y_ indicates the plasticity of material under compression. Smaller the value of 1/K, higher the plasticity of material. Using the value of intercept ‘A’ densification of powder at different stages (D_A_ and D_B_) was calculated Odeku and colleagues (*14*, Mustapha et al. (*19*, Mohammed et al. (*20*).

##### Method of applying Kawakita equation

Using Kawakita equation 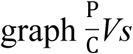. P is plotted and values of constants (a and b) are determined from slope and intercept of the plot. Reciprocal of slope is the value of ‘a’ (i.e. a = 1/slope) and reciprocal of the intercept yields ‘ab’ (i.e. ab = 1/intercept). So ‘b’ is calculated as:

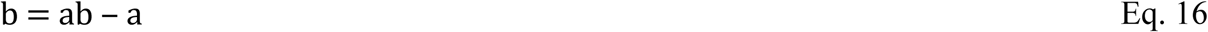

Reciprocal of ‘b’ is a pressure (P_K_) which reduces the thickness of powder bed by 50%.

#### Evaluation of In-Vitro Release behavior

Six tablets of each test (FA-FH) and immediate release marketed (reference) formulations were placed in dissolution apparatus (USP Apparatus-II: Paddle Stirring Element) containing 900 mL of dissolution media at 37± 0.5°C and 50 rpm.. Comparison was conducted in different dissolution media i.e. 0.1N HCl, phosphate buffer pH 4.5 and 6.8. 10 ml sample was withdrawn at 5, 10, 15, 20, 30, 45, 60, 90 and 120 min interval and substituted with fresh 10 ml of same solution. Each test and reference solutions were diluted, filtered and analyzed spectrophotometrically at 274nm Sharma and colleagues (*18*).

#### Release Kinetics

##### Model-Dependent Method

Various kinetic models were used for the evaluation of release pattern Costa and Lobo (*21*, Vudathala and Rogers (*22*) as mention in below equations:

###### First order kinetics

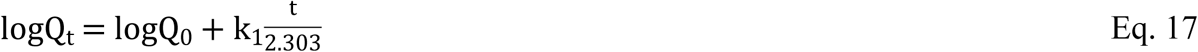

###### Weibull mode

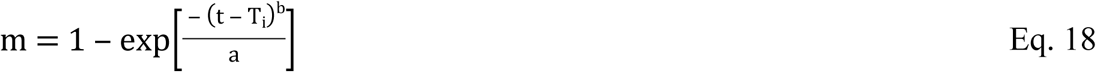

###### Hixson – Crowell modell

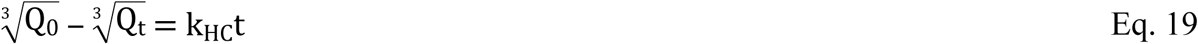

###### Higuchi model

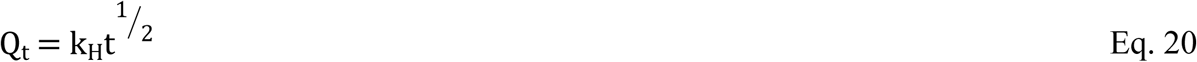

## RESULTS AND DISCUSSION

The purpose of this study was to prepare fast dispersible tablets of Aceclofenac and to evaluate the effect of different concentration of avicel PH102 on the compressional behavior of newly developed formulations. Central composite design was used for designing fast dispersible formulations and composition of fifteen fast dispersible aceclofenac formulations is given in Table 1.

### Flow Properties

Powder blends of all formulations were evaluated for true, bulk and tapped densities and results were found in the range of 1.44-1.47, 0.44-0.55 and 0.51-0.63 g/mL respectively. Flow properties of all formulations were assessed by using Hausner’s ratio, Carr’s index and angle of repose and their respective values were found to be 1.13-1.24, 11.18-19.09 % and 33.43-38.31° respectively, indicating better flow properties. Formulation blends which failed to meet the acceptable limits of micromeritics characterization were not subjected to direct compression. Since formulation blends of F13, F17 and F18 contained excessive amount of mannitol which is hygroscopic in nature, therefore, imparted negative effect on flow. Although same concentration of mannitol (25%) was present in formulations F2 and F3 but their flow was acceptable due to the presence of lesser amount of Avicel PH102. The total weight of formulations F20 was higher than the required therefore this formulations was not compressed. After micromeritic evaluation remaining formulation were subjected to compression by direct compression method using hand held hydraulic press.

### Disintegration time

All formulations except F4, F12 and F14 showed acceptable disintegration time ranging from 17-45 sec. Formulations F4 and F12 contained 0.5% and 0.1% superdisintegrant respectively therefore failed to meet the disintegration time of fast dispersible tablets. Aci-di-sol in the concentration of 1.75% indicated acceptable disintegration time but the formulation F14 having same amount of superdisintegrant presented disintegration time of 4min which is beyond the requirement of fast dispersible tablets. This difference might be due to the presence of lesser concentration of mannitol (7.63%) which is hygroscopic in nature thus facilitating the process of disintegration.

On the basis of disintegration time eight formulations were selected for further evaluation. These formulations were arranged and assigned with codes for identification purpose i.e. FA-FH, presented in Table 2. Results of micromeritic properties of selected formulation blends are shown in Table 3. Selected formulations (FA-FH) were assessed by different quality tests and results were found to be in adequate limits. Results of different physicochemical properties of compressed tablets are shown in Table 4.

**Table 2:** Composition of different fast dispersible aceclofenac (100 mg) tablet formulations used for compressional analysis.

| Formulation Codes | Ingredients Proportion (%) |  |  |  |  |  | Ingredients Amounts (mg) |  |  |  |  |  |  |  |  |
| --- | --- | --- | --- | --- | --- | --- | --- | --- | --- | --- | --- | --- | --- | --- | --- |
|  | Avicel PH102 | Mannitol | Aci-di-sol | Aspartame | Talc | Flavor | Avicel PH102 | Mannitol | Aci-di-sol | Aspartame | Talc | Flavor | Drug | Tablet weight |  |
|  | X1 (%) | X2 (%) | X3 (%) | (%) | (%) | (%) | X1 (mg) | X2 (mg) | X3 (mg) | (mg) | (mg) | (mg) | (mg) | (mg) |  |
| **FA** | 20.0 | 25.0 | 0.5 | 2% | 2% | 1% | 41 | 51 | 1.02 | 4.50 | 4.50 | 2.00 | 100 | 203.02 | 203 |
| **FB** | 20.0 | 25.0 | 3.0 | 2% | 2% | 1% | 43 | 53 | 6.50 | 4.50 | 4.50 | 3.50 | 100 | 212.53 | 213 |
| **FC** | 27.5 | 17.5 | 1.75 | 2% | 2% | 1% | 57 | 37 | 3.64 | 4.50 | 4.50 | 2.50 | 100 | 207.69 | 208 |
| **FD** | 27.5 | 17.5 | 3.39 | 2% | 2% | 1% | 59 | 38 | 7.10 | 4.25 | 4.25 | 2.50 | 100 | 214.16 | 214 |
| **FE** | 35.0 | 10.0 | 0.5 | 2% | 2% | 1% | 72 | 20 | 1.02 | 4.50 | 4.50 | 2.00 | 100 | 203.04 | 203 |
| **FF** | 35.0 | 10.0 | 3.0 | 2% | 2% | 1% | 75 | 21.5 | 7.00 | 4.50 | 4.50 | 2.50 | 100 | 212.38 | 212 |
| **FG** | 20.0 | 10.0 | 0.5 | 2% | 2% | 1% | 31 | 15 | 0.78 | 3.00 | 3.00 | 1.50 | 100 | 155.44 | 155 |
| **FH** | 20.0 | 10.0 | 3.0 | 2% | 2% | 1% | 32 | 16 | 4.83 | 3.00 | 3.00 | 1.00 | 100 | 161.18 | 161 |

**Table 3:**
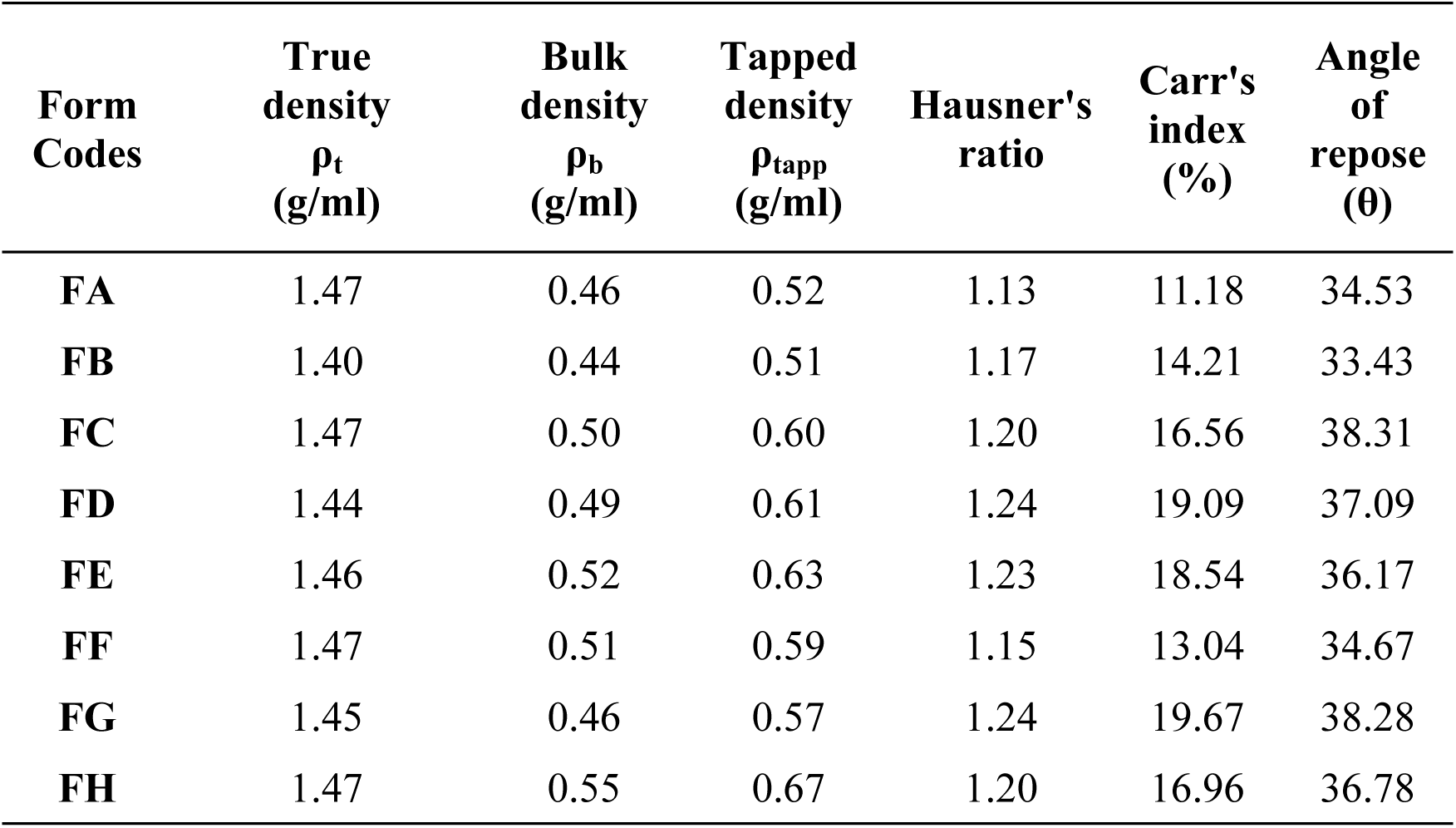
Micromeritic properties of different aceclofenac fast dispersible tablet formulations (FA-FH)

| **Form Codes** | **True density**<br>**$\rho_t$**<br>**(g/ml)** | **Bulk density**<br>**$\rho_b$**<br>**(g/ml)** | **Tapped density**<br>**$\rho_{tapp}$**<br>**(g/ml)** | **Hausner's ratio** | **Carr's index (%)** | **Angle of repose ($\theta$)** |
| --- | --- | --- | --- | --- | --- | --- |
| **FA** | 1.47 | 0.46 | 0.52 | 1.13 | 11.18 | 34.53 |
| **FB** | 1.40 | 0.44 | 0.51 | 1.17 | 14.21 | 33.43 |
| **FC** | 1.47 | 0.50 | 0.60 | 1.20 | 16.56 | 38.31 |
| **FD** | 1.44 | 0.49 | 0.61 | 1.24 | 19.09 | 37.09 |
| **FE** | 1.46 | 0.52 | 0.63 | 1.23 | 18.54 | 36.17 |
| **FF** | 1.47 | 0.51 | 0.59 | 1.15 | 13.04 | 34.67 |
| **FG** | 1.45 | 0.46 | 0.57 | 1.24 | 19.67 | 38.28 |
| **FH** | 1.47 | 0.55 | 0.67 | 1.20 | 16.96 | 36.78 |

**Table 4:** Quality attributes of different aceclofenac fast dispersible tablet formulations at different compressional pressures.

| Test parameters | Formulation Codes |  |  |  |  |  |  |  |
| --- | --- | --- | --- | --- | --- | --- | --- | --- |
|  | FA | FB | FC | FD | FE | FF | FG | FH |
| Weight (mg) | 204.1 ± 0.00 | 212.1 ± 0.00 | 207.4 ± 0.00 | 214.2 ± 0.00 | 204.1 ± 0.00 | 212.2 ± 0.00 | 155.1 ± 0.00 | 161.3 ± 0.00 |
| Diameter (mm) | 8.46 ± 0.00 | 8.48 ± 0.00 | 8.49 ± 0.00 | 8.47 ± 0.00 | 8.47 ± 0.00 | 8.48 ± 0.00 | 8.45 ± 0.00 | 8.47 ± 0.00 |
| Thickness (mm) | 2.72 ± 0.00 | 2.76 ± 0.00 | 2.72 ± 0.00 | 2.77 ± 0.00 | 2.75 ± 0.00 | 2.73 ± 0.00 | 2.66 ± 0.00 | 2.73 ± 0.00 |
| Hardness (N) | 35.59 ± 0.18 | 37.75 ± 0.14 | 40.40±0.26 | 41.38 ± 0.27 | 50.99±0.23 | 47.26 ± 0.23 | 38.14 ± 0.13 | 36.08 ± 0.12 |
| Tensile strength (MN/m<sup>2</sup>) | 5.24 | 5.67 | 6.01 | 6.19 | 7.61 | 7.05 | 4.69 | 4.95 |
| Relative density (g/cm<sup>3</sup>) | 0.93 ± 0.01 | 0.95 ± 0.01 | 0.92± 0.02 | 0.93±0.02 | 0.92 ± 0.03 | 0.92 ± 0.02 | 0.70 ± 0.03 | 0.73 ± 0.03 |
| Friability (%) | 0.5 | 0.68 | 0.30 | 0.32 | 0.34 | 0.29 | 0.76 | 0.65 |
| Disintegration time (sec) | 18 | 21 | 40 | 36 | 45 | 40 | 17 | 27 |
| Assay (%) | 99.95 ± 1.34 | 100.10 ± 0.68 | 99.63 ± 0.48 | 100.58 ± 1.68 | 98.63 ± 0.80 | 99.56 ± 1.23 | 99.64 ± 0.62 | 99.12 ± 0.61 |
| Dissolution (%) | 99.42 ± 0.52 | 100.38 ± 0.71 | 99.05 ± 0.90 | 100.35 ± 0.77 | 99.10 ± 1.04 | 100.10 ± 1.44 | 102.47 ± 0.58 | 100.34 ± 0.83 |

### RSM plot and ANOVA Summary

It is indicated in RSM plot that disintegration time was increased with increased concentration of avicel PH102 and mannitol, shown in Figure 1a and 1b. The ANOVA summary for the first response (disintegration) indicated that the model F value was 4.54 and the Probability value was less than 0.05 indicating that the quadratic model was significant. The “Adeq Precision” was 8.196 which indicated adequate signal-to-noise ratio and the design space could be navigated by the model. RSM plot Figure 1c and 1d indicated that friability was decreased with increased concentration of avicel PH 102 and mannitol. For the second response friability, F value, Probability and Adeq Precision were 10.18, < 0.05 and 11.265 respectively indicating quadratic model was valuable with the satisfactory signal. RSM plot Figure 1e and 1f presented that hardness of fast dispersible aceclofenac tablets was increased with higher concentration of avicel PH102 and mannitol. F value, Probability and Adeq Precision for third response hardness were 10.18, < 0.05 and 11.265 respectively showing model terms and linear model were acceptable with adequate signal. If A = Avicel PH102, B = Mannitol and C = Aci-Di-Sol then the final equations in terms of coded factors for disintegration, friability and hardness are given below:

**Figure 1:**
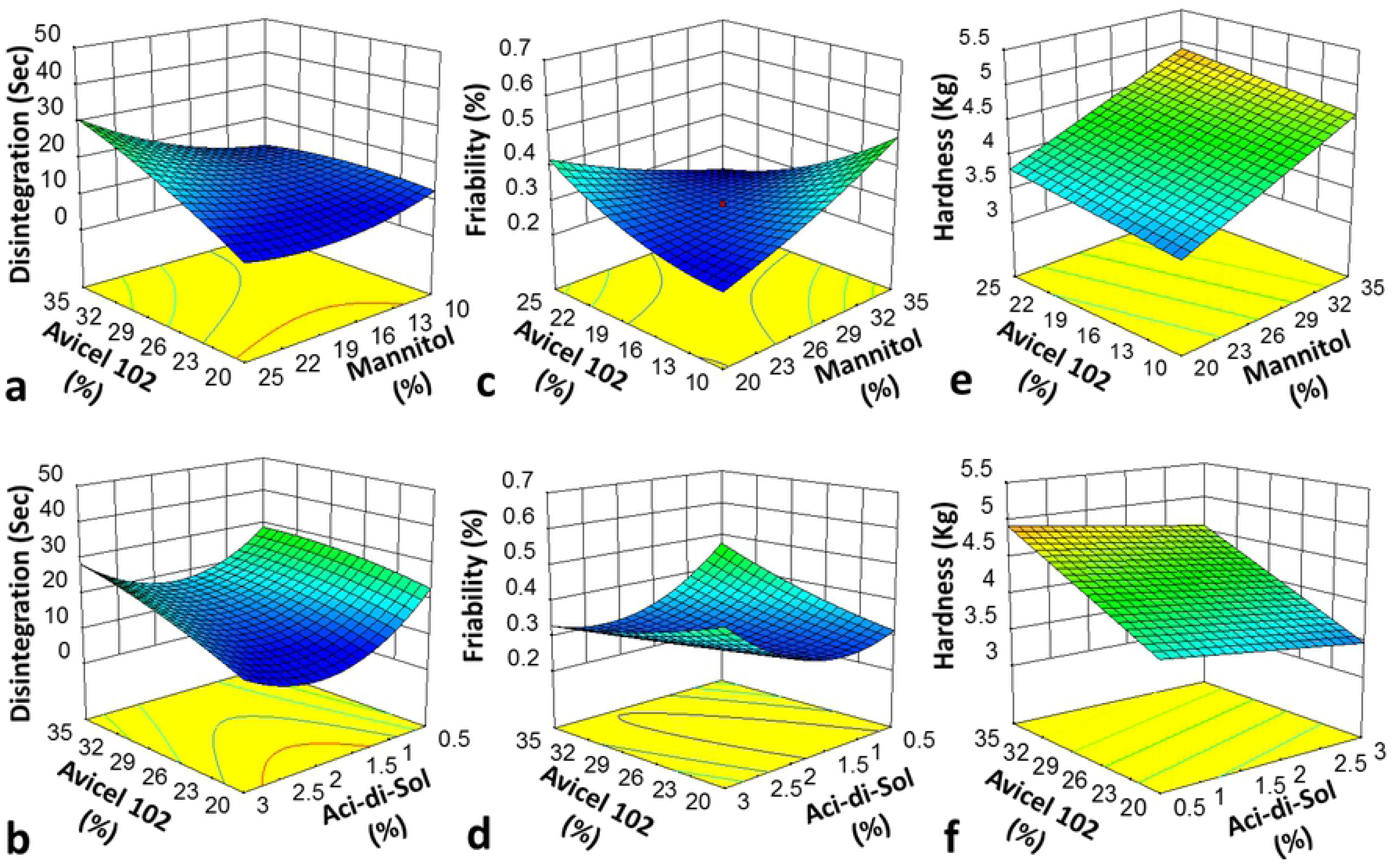
3D Response surface plots of different fast dispersible Aceclofenac tablet formulations presenting effect of independent variables on (a & b) disintegration time, (c & d) Friability and (e & f) Hardness

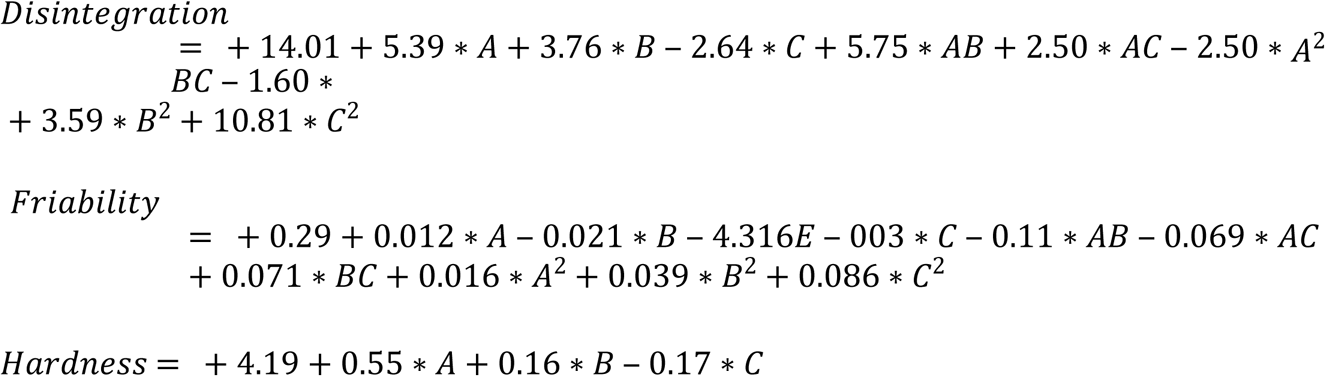

### Tensile strength and hardness

Mechanical strength is another important parameter which represents inter-particulate bonding. For tablet dosage form it is recommended to determine the mechanical properties of dominant ingredient so as to predict the overall compressional behaviour of tablets Amin et al. (*23*). Mechanical properties were estimated by measuring the tensile strength which was in the range of 4.69 - 7.61 MN/m^2^ Tablet hardness was found to be in the range from 35.59 ± 0.18-50.99 ± 0.23N. Results of tensile strength and hardness indicated sufficient mechanical strength of all formulations due to the development of strong inter-particle bonds of powder blends and % friability of the tablets was 0.29-0.76%.

### Compressional behavior analysis

For evaluating compressional behavior, ten tablets from each formulation were prepared by applying different compressional pressure 7.72, 23.16, 30.88, 38.0, 46.32, 54.04, 61.76, 69.48 and 77.2 MN/m^2^. Heckel and Kawakita equations were employed for estimating the compressional behavior of fast dispersible Aceclofenac tablets prepared by Avicel PH102. Previously it was reported that both these compaction equations are suitable for describing the compression process of powder materials based on Avicel PH 102 (*24*). Different parameters of Heckel and Kawakita equations were derived from these plots i.e. D_o,_ D_A,_ D_B,_ P_Y_, Dl and P_K_ and reported in Table 5. Heckel and Kawakita plots are presented in Figure 2 and 3 respectively.

**Table 5:** Compressional parameters obtained from heckel and kawakita equations of formulation blends.

| Formulation Codes | Heckel Parameters |  |  |  | Kawakita Parameters |  |
| --- | --- | --- | --- | --- | --- | --- |
| | $D_0$ | $D_A$ | $D_B$ | $P_Y$ (MN/m<sup>2</sup>) | $DI = 1-a$ | $P_k = 1/b$ (MN/m) |
| **FA** | 0.313 | 0.892 | 0.579 | 86.950 | 0.325 | 1.712 |
| **FB** | 0.354 | 0.922 | 0.568 | 89.520 | 0.369 | 1.040 |
| **FC** | 0.343 | 0.870 | 0.527 | 75.750 | 0.357 | 4.180 |
| **FD** | 0.340 | 0.883 | 0.543 | 74.930 | 0.349 | 4.350 |
| **FE** | 0.355 | 0.869 | 0.514 | 66.660 | 0.367 | 6.560 |
| **FF** | 0.339 | 0.863 | 0.524 | 67.560 | 0.350 | 7.690 |
| **FG** | 0.310 | 0.624 | 0.314 | 178.570 | 0.396 | -2.680 |
| **FH** | 0.370 | 0.671 | 0.301 | 212.760 | 0.474 | -3.320 |

**Figure 2:**
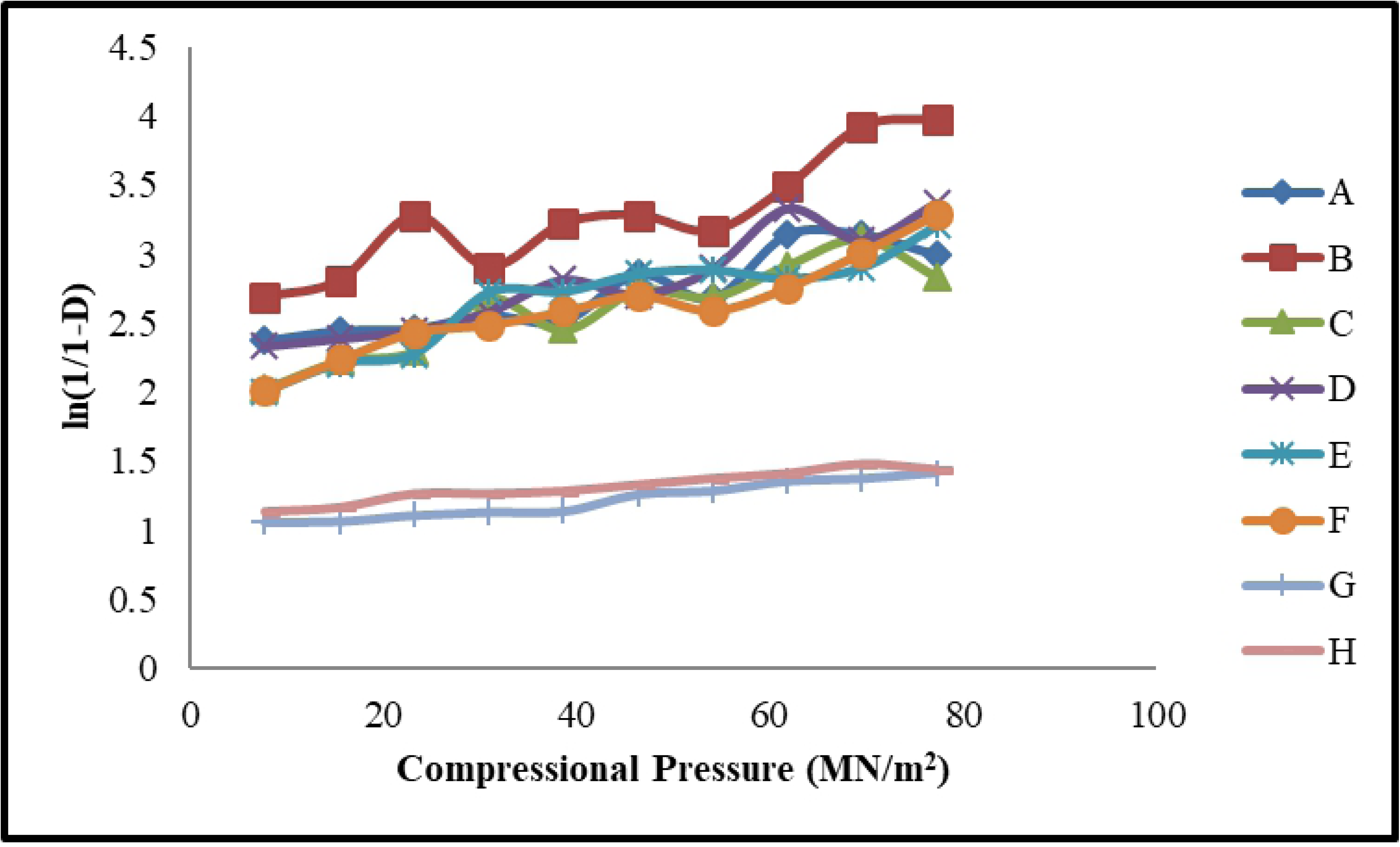
Heckel plots of fast dispersible Aceclofenac tablets.

**Figure 3:**
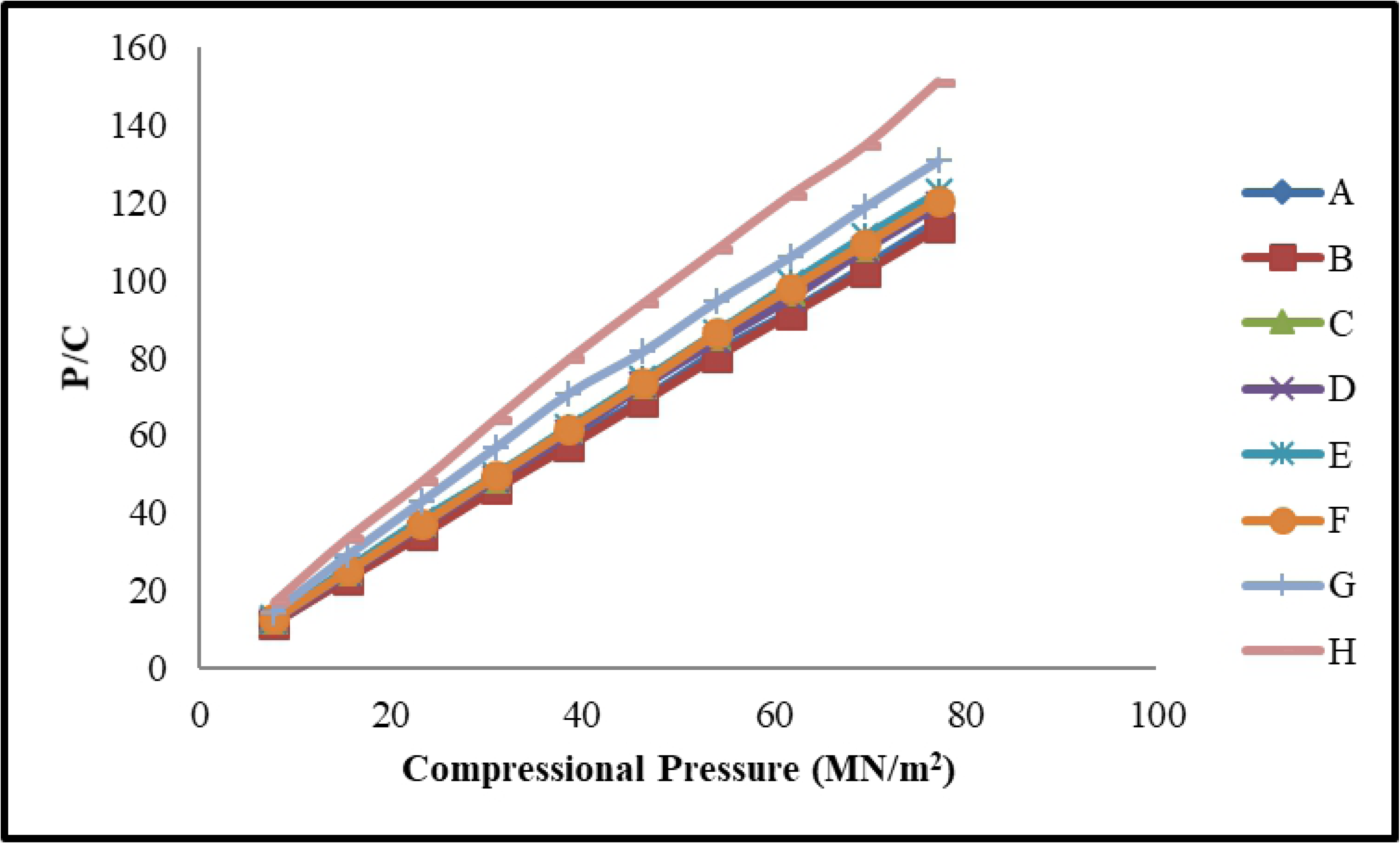
Kawakita plots of fast dispersible Aceclofenac tablets.

Heckel and Kawakita equations were used to study the compressional behaviour of all formulation. The parameters of Heckel and Kawakita equations were derived from their plots i.e. DO, D_A_, D_B_, P_Y_, D_I_ and P_K_ as mentioned in Table 4 and Figures 2 and 3. It was reported that DO values for different aceclofenac formulations were increased as the concentration of binder was increased. This phenomenon indicates that at die filling the initial packing of the formulations increases with the higher concentration of binder Amin and colleagues (*23*). In formulations FC, FD, FE and FF, the amount of avicel PH 102 was high (27.5 – 35 %) which resulted in increased hardness, tensile strength and less percentage friability (Table 3). In another study conducted by Odeku *et al.*, in 2005, it was reported that the value of DO was higher with increased concentration of starch (as binder) in Paracetamol tablets Amin and colleagues (*23*). Scientists also found that compaction behaviour of powder bed mostly dependent upon the deformation properties of the blends and the applied processing methods Amin and colleagues (*23*).

### Heckel analysis

From Heckel plots it was found that the initial packing of powder blends (D_o_) in FE was found to be high i.e. D_o_ = 0.355 as FE contained increased concentration of avicel PH102 (35 %). The values of D_B_ and D_A_ were calculated from Eq. 2 and 3 respectively. The D_A_ values at zero and low pressures presented the total degree of packing. Generally formulations with increased concentration of avicel 102 exhibited the lowest values of D_A_. The D_A_ value which showed the degree of packing at low compressional pressure was found lowest in FE (0.869) and FF (0.863). Values of D_A_ for different formulations observed in the presented sequence: FF<FE<FC<FD<FA<FB. Formulation FG and FH were found to be outliers Amin and colleagues (*23*).

The D_B_ values presented the densification of powder bed at low pressure which shows the particles rearrangement by applying compressional pressure leading to the particle fragmentation. This fragmentation could be plastic or elastic. It was observed that formulations containing high amount of avicel PH102 i.e. FA (0.568) and FB (0.579) showed decreased values of D_B_. The sequence is FE<FF<FC<FD<FA<FB. Amin *et al*., 2012 stated that each powder has its own compressional characteristics. In the form of powder blend these characteristics get significantly changed which affects the tablet stability Amin and colleagues (*23*). Formulations FG and FH were found to be outliers for D_A_ values which could be due to formualtion blend. It was observed that all formulations yielded low values of D_o_ than D_B_ due to particles fragmentation and particles rearrangement in the die at reduced pressure. Generally, increased porosity of powder blend at zero pressure yields low values of D_o_ Amin and colleagues (*23*).

Another parameter ‘P_Y_’ (mean yield pressure) is the measure of plasticity of the material. A plastic material is desirable for compression while elastic material creates problem during compression due to elastic recovery. Formulations having greater tendency to deform plastically usually have low values of P_Y_. In this study FE (66.66 MN/m^2^) and FF (67.56 MN/m^2^) showed the lowest ‘P_Y_’ value and the highest plasticity due to the presence of highest amount of avicel PH102, whereas, formulations FG and FH showed the lowest plasticity when compressional pressure was applied (Table 5). It means that higher concentration of avicel PH102 rendered the tablets more plastic and thus tablet manufacturing process became easy.

### Kawakita analysis

The Kawakita plot was also constructed to evaluate the compressional behavior of formulations (Figure 2). Kawakita plots presented a linear relationship at different compressional pressures with correlation coefficient above 99% for formulations FA-FH. From the slope and intercept of Kawakita plots the values of *a* and *ab* were determined respectively. Values of 1-*a* indicated initial relative density (D_I_) of the formulations. By using reciprocal of *b* values, the inverse measurement of plasticity (P_k_) was estimated as given in Table 5. It was observed that initial relative density of formulations (D_I_) was decreased in formulations containing higher concentration of avicel PH102. The initial relative density (D_I_) of all formulations was in the range of 0.35-0.47. Formulations containing higher concentration of avicel PH102 showed low D_I_ value. Results indicated that higher D_I_ values were observed as compared to the corresponding values of DO, which has been previously reported by other researches Amin and colleagues (*23*). On the other hand the P_K_ values of these formulations were high.

It was observed in the present study that as concentration of avicel PH102 was increased the P_K_ value was also increased (1.04 - 7.69 MN/m) Formulations FE and FF had the highest P_K_ values and FA and FB had the lowest P_K_ values. The results of this parameter showed the elastic nature of avicel PH102 which is predominant among other excipients in the formulations. It was found from Heckel and Kawakita parameters analysis that FE and FF showed the fastest onset of plastic deformation whereas FA and FB showed maximum plastic deformation in combination with aceclofenac and other excipients. However no clear cut variation pattern of Heckel and Kawakita parameters was observed as indicated in Table 5.

### In Vitro Dissolution

In the present study *in vitro* drug release profiles of newly developed and optimized aceclofenac formulations were determined in different dissolution media as shown in Figure 4 (A-C). All formulations demonstrated maximum dug release in phosphate buffer pH 6.8. Different kinetic models were used to analyze the release behavior of formulations FA-FH. Results indicated that all formulations followed First-order and Weibull model in different dissolution media with highest r^2^ values found in phosphate buffer pH 6.8 i.e. 0.946 - 0.954 and 0.986 - 0.996 respectively as shown in Table 6.

**Table 6:** Release kinetics of fast dispersible aceclofenac (100 mg) tablets at different pH.

| Formulation Codes | First Order |  | Higuchi |  | Hixson-Crowell |  | Weibull model |  |  |
| --- | --- | --- | --- | --- | --- | --- | --- | --- | --- |
| | $r^2$ | $\frac{K}{(\text{hr}^{-1})}$ | $r^2$ | $\frac{K_H}{(\text{hr}^{-1/2})}$ | $r^2$ | $\frac{K_{HC}}{(\text{hr}^{-1/3})}$ | $r^2$ | $\alpha$ | B |
| **0.1N HCl** |  |  |  |  |  |  |  |  |  |
| **Reference** | 0.374 | 0.007 | 0.765 | 4.622 | 0.645 | 0.001 | 0.893 | 9.695 | 0.378 |
| **FA** | 0.353 | 0.009 | 0.768 | 5.046 | 0.664 | 0.001 | 0.963 | 8.065 | 0.367 |
| **FB** | 0.408 | 0.009 | 0.796 | 5.107 | 0.696 | 0.002 | 0.939 | 8.181 | 0.379 |
| **FC** | 0.323 | 0.009 | 0.756 | 5.043 | 0.653 | 0.001 | 0.925 | 7.764 | 0.359 |
| **FD** | 0.349 | 0.009 | 0.760 | 5.106 | 0.657 | 0.001 | 0.925 | 7.812 | 0.364 |
| **FE** | 0.324 | 0.009 | 0.751 | 5.031 | 0.646 | 0.001 | 0.919 | 7.803 | 0.359 |
| **FF** | 0.330 | 0.009 | 0.760 | 5.089 | 0.659 | 0.001 | 0.925 | 7.745 | 0.362 |
| **FG** | 0.311 | 0.009 | 0.744 | 5.052 | 0.641 | 0.001 | 0.916 | 7.673 | 0.357 |
| **FH** | 0.346 | 0.009 | 0.779 | 5.044 | 0.680 | 0.001 | 0.932 | 7.985 | 0.366 |
| **pH 4.5** |  |  |  |  |  |  |  |  |  |
| **Reference** | 0.681 | 0.041 | 0.868 | 6.480 | 0.809 | 0.002 | 0.950 | 9.179 | 0.483 |
| **FA** | 0.948 | 0.021 | 0.943 | 8.357 | 0.944 | 0.004 | 0.985 | 17.192 | 0.733 |
| **FB** | 0.946 | 0.022 | 0.945 | 8.418 | 0.950 | 0.004 | 0.988 | 17.079 | 0.736 |
| **FC** | 0.950 | 0.022 | 0.950 | 8.391 | 0.951 | 0.004 | 0.991 | 16.625 | 0.726 |
| **FD** | 0.954 | 0.022 | 0.946 | 8.474 | 0.950 | 0.004 | 0.989 | 17.511 | 0.744 |
| **FE** | 0.964 | 0.022 | 0.953 | 8.527 | 0.958 | 0.005 | 0.987 | 20.903 | 0.785 |
| **FF** | 0.950 | 0.021 | 0.943 | 8.359 | 0.941 | 0.004 | 0.982 | 17.076 | 0.730 |
| **FG** | 0.953 | 0.022 | 0.939 | 8.453 | 0.944 | 0.004 | 0.984 | 17.538 | 0.743 |
| **FH** | 0.952 | 0.021 | 0.942 | 8.401 | 0.940 | 0.004 | 0.987 | 15.959 | 0.717 |
| **pH 6.8** |  |  |  |  |  |  |  |  |  |
| **Reference** | 0.948 | 0.044 | 0.781 | 11.553 | 0.969 | 0.012 | 0.996 | 41.658 | 1.267 |
| **FA** | 0.946 | 0.051 | 0.736 | 11.438 | 0.950 | 0.012 | 0.991 | 27.069 | 1.223 |
| **FB** | 0.954 | 0.052 | 0.733 | 11.339 | 0.951 | 0.012 | 0.992 | 21.981 | 1.144 |
| **FC** | 0.948 | 0.051 | 0.713 | 11.351 | 0.944 | 0.012 | 0.996 | 29.015 | 1.248 |
| **FD** | 0.952 | 0.051 | 0.741 | 11.398 | 0.953 | 0.012 | 0.993 | 21.695 | 1.111 |
| **FE** | 0.951 | 0.050 | 0.728 | 11.240 | 0.948 | 0.012 | 0.992 | 20.114 | 1.080 |
| **FF** | 0.949 | 0.050 | 0.735 | 11.374 | 0.950 | 0.012 | 0.989 | 24.857 | 1.165 |
| **FG** | 0.949 | 0.050 | 0.729 | 11.397 | 0.949 | 0.012 | 0.995 | 28.654 | 1.229 |
| **FH** | 0.947 | 0.050 | 0.739 | 11.357 | 0.949 | 0.012 | 0.986 | 21.034 | 1.088 |

**Figure 4:**
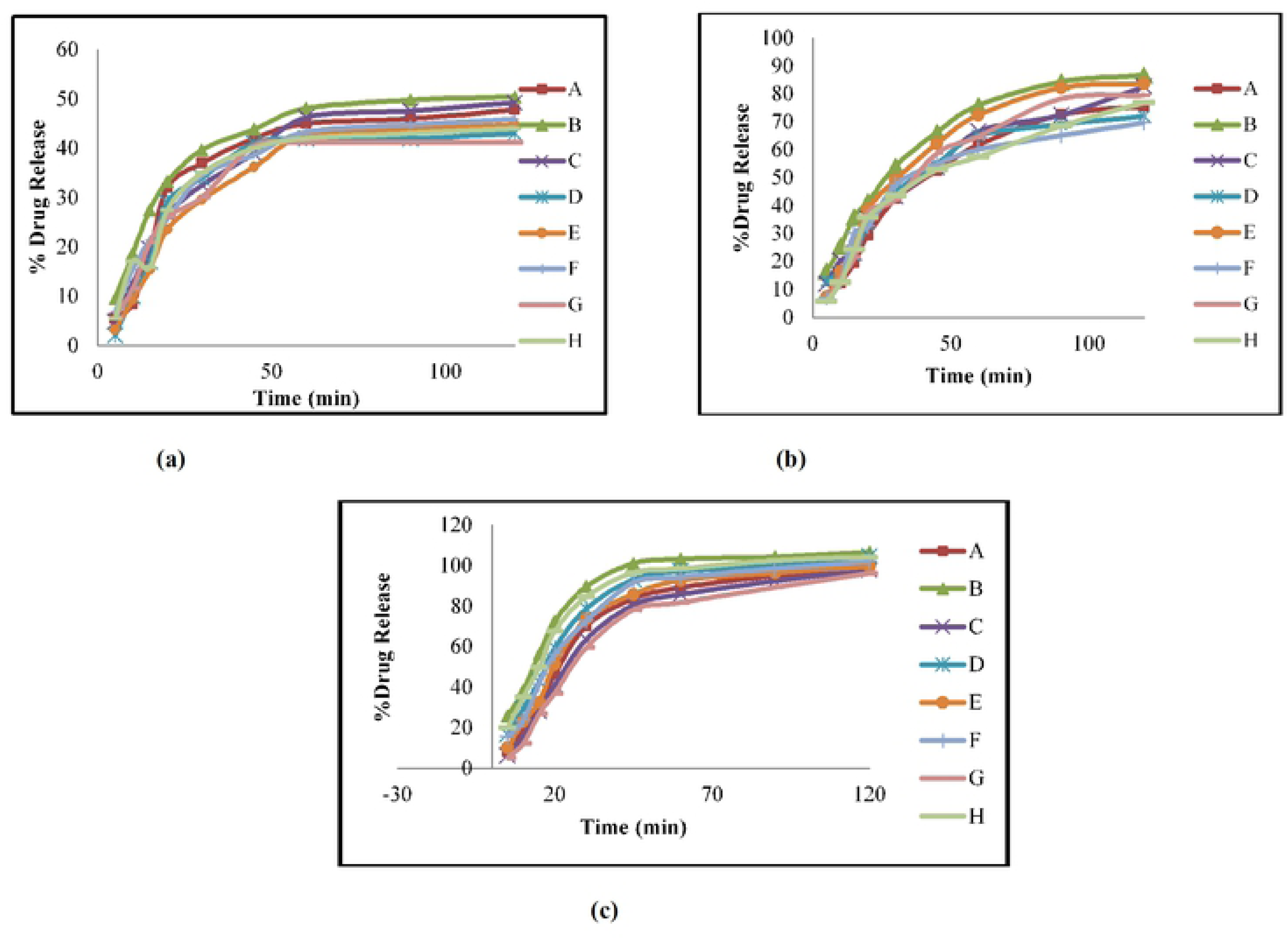
Drug release kinetics of Aceclofenac fast dispersible tablets in 900 ml of (a) pH 1.2 (b) phosphate buffer pH 4.5 (c) phosphate buffer pH 6.8

## CONCLUSION

In this study fast dispersible aceclofenac tablets were prepared and the effect of avicel PH102 was examined on compressional, mechanical and release properties of fast dispersible aceclofenac formulations. Using the Heckel and Kawakita equations, the compressional behavior was observed. The concentration of avicel PH102 exhibited a significant impact on the compressional, mechanical and release properties of the Aceclofenac fast dispersible formulations. Hence a suitable selection of excipient with appropriate concentration is important at formulation development stage to ensure stable, elegant and palatable dosage form for the patient.

## List of Tables

Table 1: Composition of fast dispersible aceclofenac formulations using central composite design

Table 2: Composition of different fast dispersible aceclofenac (100 mg) tablet formulations used for compressional analysis

Table 3: Micromeritic properties of different aceclofenac fast dispersible tablet formulations Table 4: Quality attributes of different aceclofenac fast dispersible tablet formulations at different compressional pressures

Table 5: Compressional parameters obtained from heckel and Kawakita equations of formulation blend

Table 6: Release kinetics of fast dispersible aceclofenac (100 mg) tablets in different pH

